# ‘Trans-differentiation of neutrophils from plasmablast’ is an artefact caused by over-reliance on machine algorithms in single cell RNA sequencing analysis: Lesson learnt and steps ahead

**DOI:** 10.1101/2025.02.07.636761

**Authors:** Jinghan Huang, Phillip S.C. Yam, Nelson L.S. Tang

**Affiliations:** Department of Chemical Pathology, Faculty of Medicine, The Chinese University of Hong Kong, Hong Kong, Hong Kong SAR, China; Department of Statistics, Faculty of Science, The Chinese University of Hong Kong, Hong Kong, Hong Kong SAR, China; Cytomics Limited, Hong Kong Science Park, Hong Kong, Hong Kong SAR, China; Li Ka Shing Institute of Health Sciences and CAS Center for Excellence in Animal Evolution and Genetics, Faculty of Medicine, The Chinese University of Hong Kong, Hong Kong, Hong Kong SAR, China

## Abstract

Single cell RNA sequencing (scRNA-seq) provides new opportunities to characterize gene expression for individual cells. However, the sparse nature of the scRNA-seq data with many zero counts or missing values presents a major challenge to its analysis. The presence of low-quality cells further complicates the analysis. Here, we showed that the trans-differentiation of plasmablasts (activated plasma cells) into neutrophils reported in COVID patients (Wilk et al., 2020 in Nature Medicine) was an artefact of trajectory analysis. It was caused by ∽30 low-quality cells linking the 2 cell populations that are of unrelated lineages in hematopoietic differentiation.

Such artefacts are not readily spotted during the current practice of peer reviews as the current statistical guidelines of most journals are not catered for big data such as that of scRNA-seq. New standards of statistics and quality control measures for machine algorithms are not in place and they are urgently needed to safeguard against over-interpretation of high dimensional data. We propose a comprehensive framework to ensure reproducibility in high-dimensional data analysis, emphasizing quality checks, sensitivity analyses, alternative and multiple algorithms validation. Finally, and most importantly, a hypothesis-driven research approach should be upheld.

## Introduction

The paper of Wilk et al. published in the July 2020 issue of Nature Medicine represented the first single-cell RNA (scRNA) sequencing of PBMC in COVID patients^1^. This has become a precious resource for the research community and we all applaud their contribution. However, about the potential of trans-differentiation of developing neutrophils from plasmablasts (PB) which was described in abstract, main text and Figure 4 of the paper (**Figure 1** in this paper), we and others^2^ expressed concerns.

**Figure 1.**
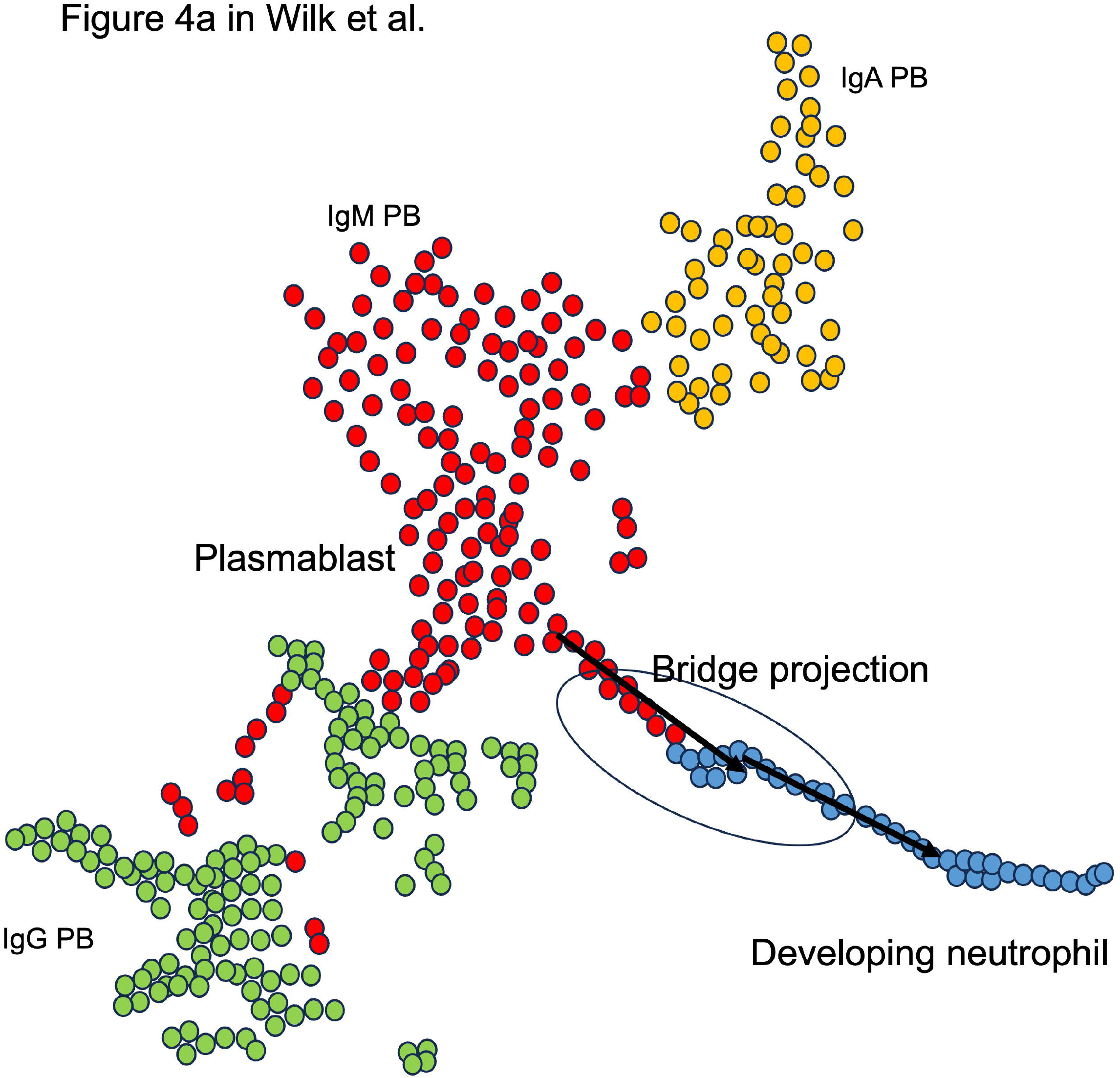
Hypothesis of trans-differentiation of IgM plasmablast to developing neutrophils in the original paper. An artwork illustration drawing to recapture the projection of trajectory analysis presented by Wilk et al. in Figure 4 of their paper.

In 2016, high ranked journals like Nature and Science spearheaded a movement against the Reproducibility Crisis in Science and proposed various study and data analysis guidelines for improvement of research studies. With the advance in machine algorithms and AI, we are afraid that current guidelines are not keeping pace with analysis of high dimensional data and new guidelines need to be developed to avoid the next outbreak of Reproducibility Crisis in Science at this era of AI.

## Methods

The processed dataset was downloaded from COVID-19 Cell Atlas (https://www.covid19cellatlas.org/#wilk20). There are 26,361 rows (genes) and 44,721 columns (cells) in this data matrix. The data matrix is sparse, with 95.67% of values are zero counts.

### Normalization methods

As for the discovery dataset GSE150728 COVID-19, conventional normalizations for scRNA-seq data were used, i.e., SC-Transform (regularized negative binomial regression; used in original study) implemented by the function SCTransform() in Seurat. Same parameters were used as applied in original study.

### Dimension reduction: PCA and UMAP

After SC-Transform, transformed data was processed by PCA using RunPCA() function, followed by running UMAP using RunUMAP() function in Seurat. The number of PCs used in UMAP processing was determined as applied in the original studies. To zoom in to the interested region, we reduced the cells to specific cell type subset and redo the PCA and UMAP for the subset. The 2-D UMAP / PCA plots were generated for all or subset cell types.

### Select potential poor-quality cells that led to bridge projection in UMAP and compare them with other cells

We hypothesized that the low-quality cells caused by dropouts may lead to unexpected patterns and suspicious findings such as incorrect trans-differentiation. Therefore, we focused on those suspicious bridge projection cells in the UMAP embeddings which was likely to generate differentiation trajectories. We used the function CellSelector() from Seurat and picked out those bridge projection cells and compared them with all other cells or other regional cells. Chi-square tests were performed across the four quartiles of number of expressed genes (nFeature_RNA) between the selected bridge projection cells and all other cells or other regional cells.

### Apply different quality control (QC) / hard filters to explore the changes of trajectory inference

Next, we investigated how these bridge projections changes and whether these suspicious trajectories would be eliminated by applying different QC standards, e.g., hard filters of different number of expressed genes (nFeature_RNA) thresholds (>200, 400, 600, etc.). The genes were ranked by their nFeature_RNA and were visualized in scatter plots. After applying each QC step, the data was re-processed by PCA and UMAP for visualization. Additionally, Monocle3 was used for predicting pseudotime information with all parameters set to default.

## Results and Discussion

The suggestion of trans-differentiation made by Wilk et al. is based on the **30 cells forming a bridge projection** which is part of the PB cluster (a projection of the PB cluster E joining to neutrophil (cluster K) by the bridge projection cluster J in Figure 4e of the paper). This feature was fed into a trajectory analysis using algorithms which inferred RNA velocity, leading to the hypothesis that developing neutrophils might trans-differentiation from plasmablasts. Subsequently, this trans-differentiation hypothesis has received a wide and far-reaching attention and has been cited in multiple articles and reviews^4–10^ and even in a monograph textbook recently^11^, despite the fact that the authors went on to disprove this hypothesis in a subsequent manuscript^12^. A senior author of the paper confirmed that this trans-differentiation could not be supported experimentally or in analysis with larger numbers of patients^12^. To the general audience unfamiliar with scRNA analysis, a clear statement to invalidate this trans-differentiation proposition is warranted to keep the record straight and avoid further perpetuation of ‘this new knowledge’ arising from algorithm artefact in trajectory analysis of scRNA seq data.

Figure 2 shows the location of this bridge formed by 30 low quality cells between PB and neutrophil in UMAP. However, they largely come from only 1 individual. Gene expression matrix of these cells is attached as an CSV file for readers for further evaluation (**Additional file 1**). **Figure 3** shows the effect of applying additional quality filter to the data and this 30-cell bridge disappeared (**Figure 3C-D**) when filtering out the lowest ∽10% of cells with least number of expressed gene (UMI count >=1). Finally, when we perform pseudotime trajectory analysis with or without these 30 low quality cells, the results support that these 30 low quality cells contribute to the artefact results of trans-differentiation of PB to neutrophils as pseudotime can no longer assigned to the remaining neutrophils after QC filter is used (**Figure 4**).

**Figure 2.**
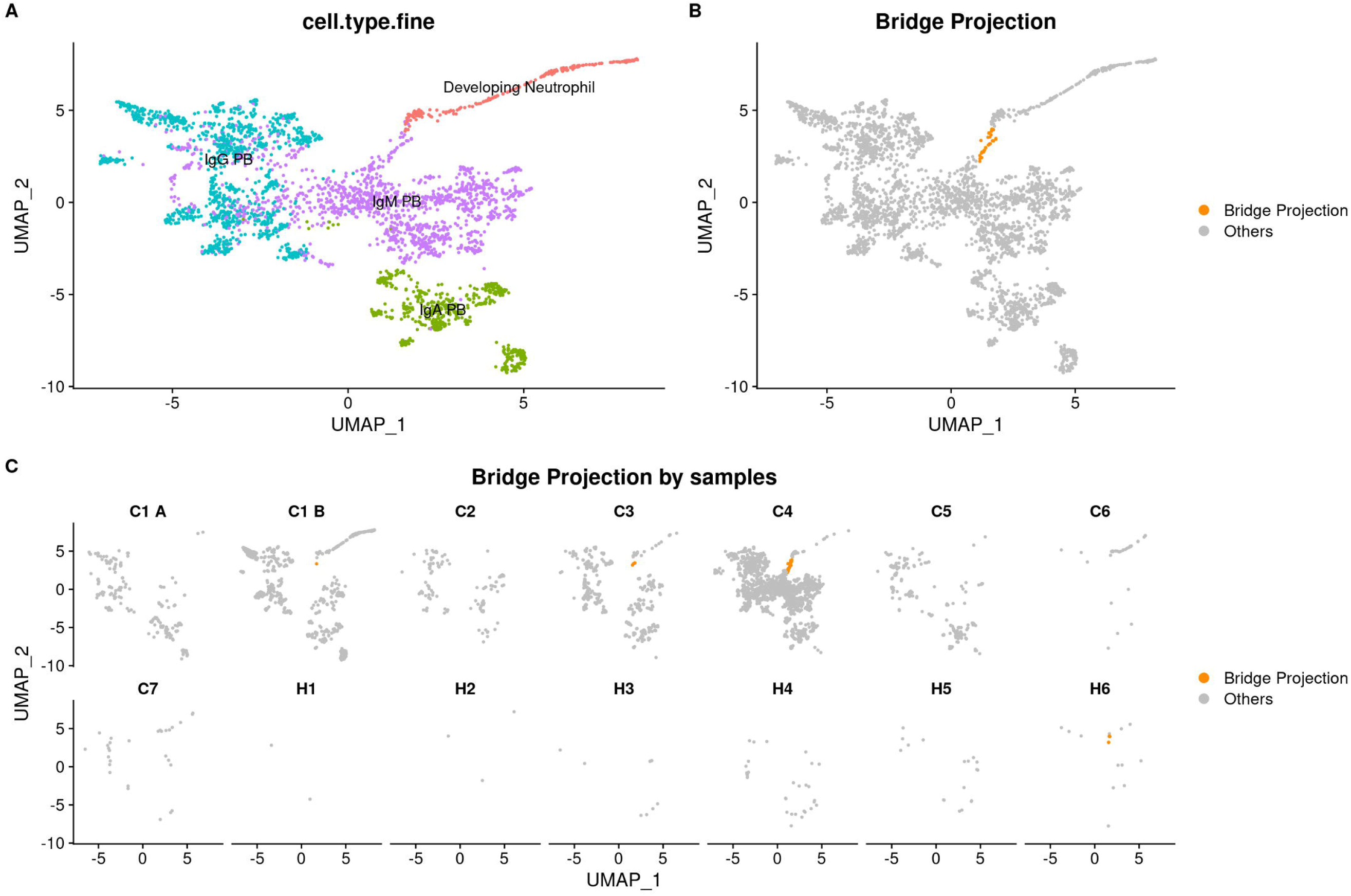
The bridge projection connecting IgM plasmablast and Neutrophils is formed by 30 cells. The bridge projection connecting IgM plasmablast and Neutrophils is formed by 30 cells and majority of them were from one patient (C4). 2A shows the clusters of PBs and neutrophils. 2B and 2C highlights the bridge projection and their origins among individual subjects.

**Figure 3.**
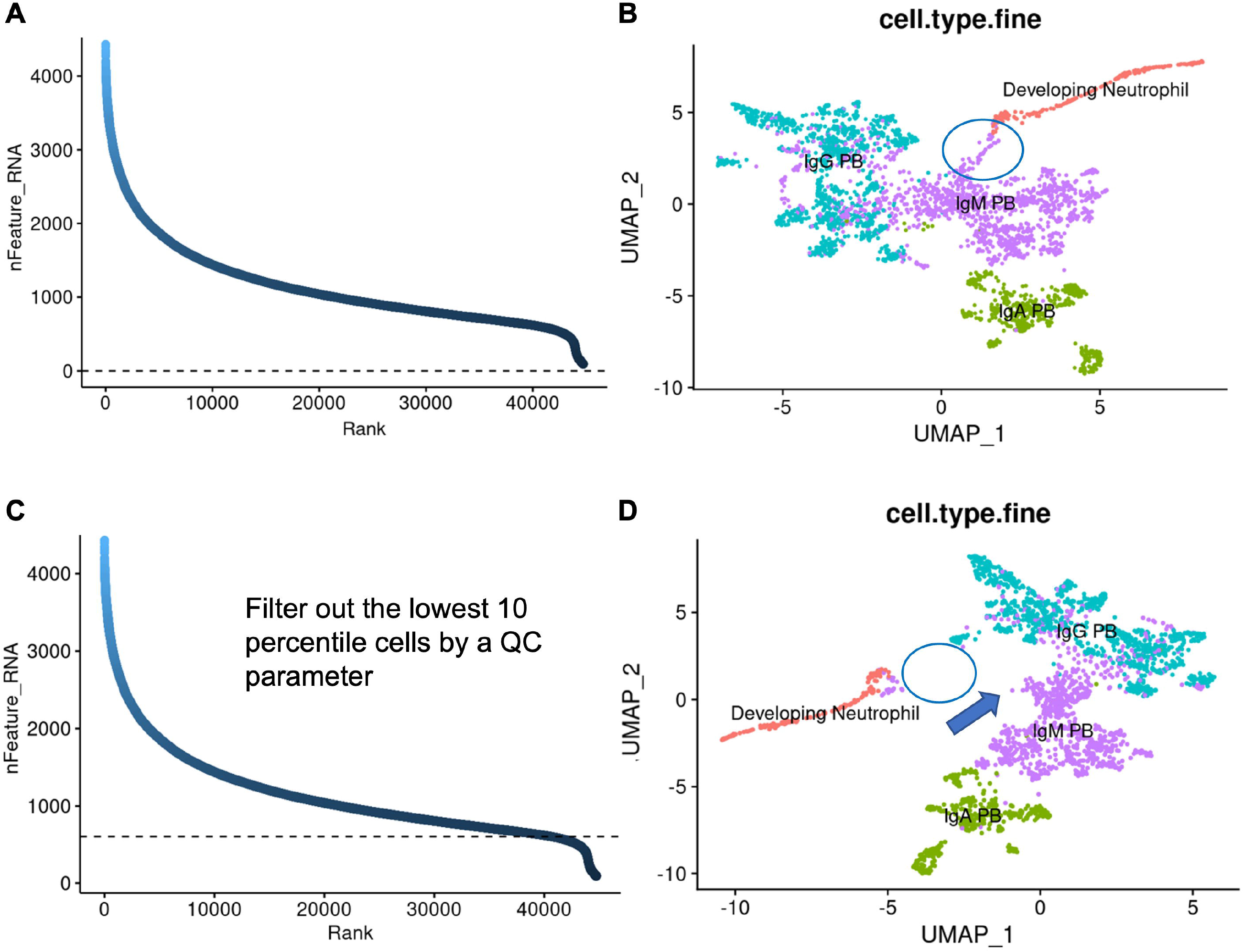
Quality of cells forming the bridge projection by nFeature (no. of gene expressed in a cell). The quality parameter nFeature (no. of genes expressed in a cell) is shown in 3A and 3C. The UMAP of all cells without QC filter is shown in 3B and the bridge projection which is part of the IgM PB cluster is marked by a blue circle. When QC filter is applied to remove the lowest 10^th^ percentile cells by nFeature (3C), the bridge projection (marked by circle) together with some adjacent IgM PB cells (marked by arrow) are filtered out (3D). This indicates that these are the low-quality cells and the bridge leading to trans-differentiation proposition (based on these low-quality cells) disappeared.

**Figure 4.**
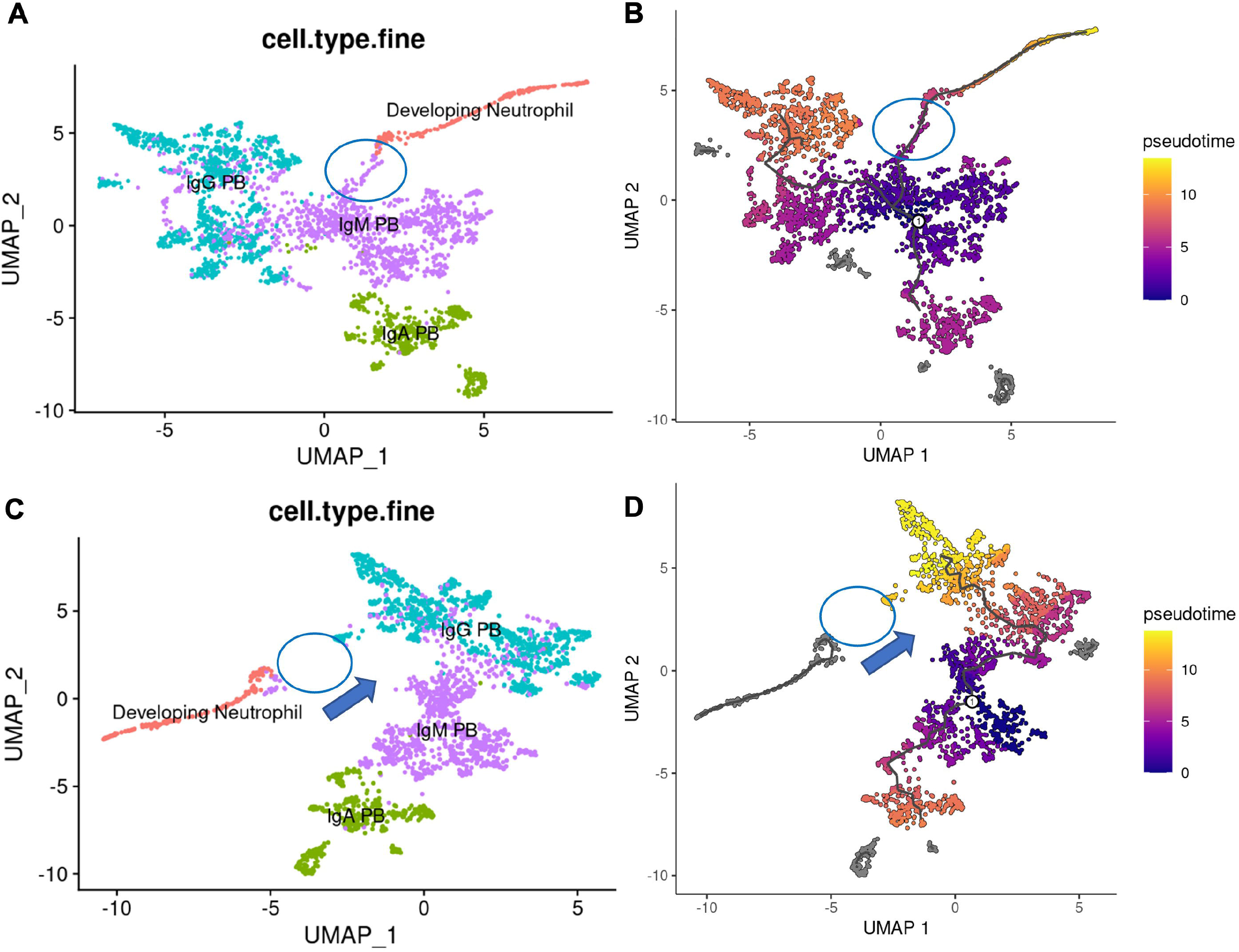
The bridge of 30 low quality cells caused trajectory algorithms (Monocle3) to report the artefact trans-differentiation of neutrophil from PB. Figure 4A-B are results of UMAP cell type clusters and pseudotime trajectory analysis when no additional filter is applied to the scRNA seq data (as in Figure 3A-B). Figure 4C-D are results of UMAP and trajectory analysis when the lowest 10^th^ percentile cells of nFeature are filtered out (as in Figure 3C-D). The bridge projection (marked by circle) together with some adjacent IgM PB cells (marked by arrow) are filtered out. In trajectory analysis, neutrophils are labeled in grey color indicating a pseudotime cannot be assigned. These results support that these 30 low quality cells contribute to the artefact in trajectory analysis.

The attention to the 2016 movement against the Reproducibility Crisis decayed and most young researchers are not aware of such campaign a decade ago. Furthermore, previous guidelines may be outdated and they need revamp with the emergence of a vast number of machine algorithms and AI. New difficulties in upholding open science are encountered like the huge volume of data (nature of big data in modern experiments) of scRNA-seq matrix which precludes most peer-reviewers from fully understanding the details of data from which conclusions are drawn. Here, we suggest a new but simple approach for the journal editorial offices to implement to avoid similar pitfalls. We showed that simple steps would be sufficient to avoid over-interpretation due to machine algorithms or AI.

### A. Quality check of the data from which conclusion is drawn

We found that cells forming the bridge projection connecting PB and developing neutrophils (only ∽30 of them and their full expression matrix are attached in **Additional file 1**) were actually among the worst quality cells in the whole scRNA-seq matrix. **Figure 2A** shows cells forming the spurious bridge and these cells had the least nCount (number of UMIs per cell representing the overall expression) and nFeature (number of genes expressed in a cell).

Furthermore, if a quality filter of valid cells was set to nFeature>600 (at about the lowest 10th percentile value of all cells), this spurious bridge (from which the trans-differentiation process was based) disappeared.

### B. Sensitivity analysis

Besides, 20 of these 30 cells (67%) were derived from one patient (patient C4, labelled as covid_558 in **Additional file 1**). Therefore, any kind of sensitivity analysis such as jackknife resampling will be able to pick up the spurious results.

### C. Reproducibility of conclusion using algorithms using different assumptions

Of course, normalization and transformation of the scRNA-seq data also plays a key role in shaping the geometry of data points in high dimensional space. Therefore, replication using alternative normalization and transformation algorithms will be required to demonstrate that any new preposition is not solely due to data transformation.

### D. Hypothesis driven research

Last but not the least, research and results interpretation should be hypothesis driven rather than being led by results. Experimental hypothesis and biological plausible interpretation are the pillars of all sound studies. The journal should uphold these principles which are utterly important in the modern era of big data.

### Recommendations for the detailed steps

In fact, many scRNA analysis tools are descriptive in nature as the high dimensional data is usually generated from only a group of few patients. Commonly used statistics are not applicable in view of the limited sample size of patients. However, some kind of statistical analysis is needed if we want to provide inference in probability terms. Here, we proposed 2 easy steps borrowed from the analysis of GWAS data which similarly handle high dimensional data.

1. Fit expected feature counts and apply a filter to outliner cells. scRNA-seq data are well known for heterogeneity in QC. Like GWAS Quantile-Quantile (QQ) plot, dataset specific fitting is allowed by this approach. Characters of interest (e.g., the bridge projection) should be persistent both before and after a QC filter. Figure shows the effect of the number-of-feature (nFeature, no. of expressed gene) filter on the bridge projection. In this example, they disappeared when the feature (gene with counts) filter was set at 600 genes per cell.
2. A test for a uniform distribution across various QC quantiles should be performed to gauge the level of confidence of the focus of interest. It could be done by a goodness of fit test using chi-square on a 4 x 2 table. First, a QC parameter is divided into quantiles and the distribution of the QC value of those cells forming the character of interest is compared to that of adjacent clusters (regional comparison) or to that of all cells (global comparison). The P values of goodness of fit were significant indicating that those cells forming the bridge projection might be a biased finding. **Table 1-2** shows the comparison of the nFeatures of these ∽30 cells in the bridge junction with either PB and neutrophils clusters (local comparison, **Table 1**) or all cells (global comparison, **Table 2**). Both comparisons showed a high level of statistical significance. This approach should be implemented as a routine for scRNAseq analysis particularly when the inference is important. Of course, a significant difference (P-value < 0.05) in the QC values does not automatically deny the plausibility of the character of interest but it will raise the alert that additional justification is needed.
3. The same feature of interest should be persistent after applying various normalization and transformation algorithms to the data. If the feature (such as the bridge junction here) is robust, it should remain even after different pre-processing methods are applied to the data.

**Table 1.** Regional comparison of quality of cells forming the bridge junction.

| Quantile of QC parameter: no. of features (genes expressed per cell) | Q1 | Q2 | Q3 | Q4 | P-value |
| --- | --- | --- | --- | --- | --- |
| Regional (developing Neutrophils, and PB) | 797 | 798 | 796 | 796 |  |
| Junction projection (30 cells) only | 17 | 9 | 4 | 0 | $9.7 \times 10^{-5}$ |

The bridge projection cells were among the-least-expressed-genes (a feature of cell quality) cells among adjacent clusters of plasmablasts.

**Table 2.**
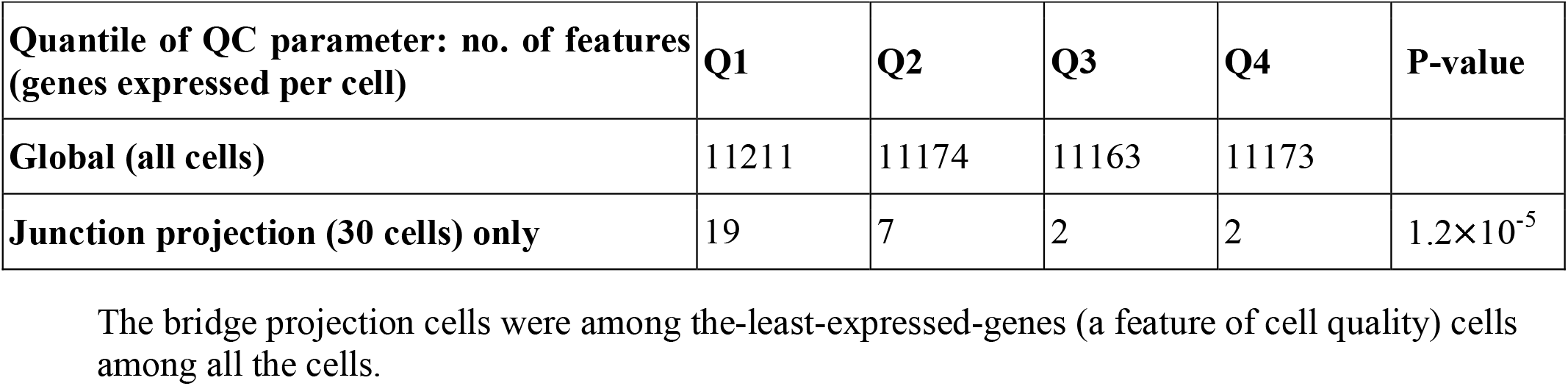
Global comparison of cell quality.

We believe that by using these statistical QC approaches, scRNAseq analysis and inference will be more streamlined in the future.

